# Metagenomics for pathogen detection during a wildlife mortality event in songbirds

**DOI:** 10.1101/2023.06.20.545358

**Authors:** Lusajo Mwakibete, Sabrina S. Greening, Katrina Kalantar, Vida Ahyong, Eman Anis, Erica A. Miller, David B. Needle, Michael Oglesbee, W. Kelley Thomas, Joseph L. Sevigny, Lawrence M. Gordon, Nicole M. Nemeth, C. Brandon Ogbunugafor, Andrea J. Ayala, Seth A. Faith, Norma Neff, Angela M. Detweiler, Tessa Baillargeon, Stacy Tanguay, Stephen D. Simpson, Lisa A. Murphy, Julie C. Ellis, Cristina M. Tato, Roderick B. Gagne

## Abstract

Mass mortality events in wildlife can be indications of an emerging infectious disease. During the spring and summer of 2021, hundreds of dead passerines were reported across the eastern US. Birds exhibited a range of clinical signs including swollen conjunctiva, ocular discharge, ataxia, and nystagmus. As part of the diagnostic investigation, high-throughput metagenomic next-generation sequencing was performed across three molecular laboratories on samples from affected birds. Many potentially pathogenic microbes were detected, with bacteria comprising the largest proportion; however, no singular agent was consistently identified, with many of the detected microbes also found in unaffected (control) birds, and thus considered to be subclinical infections. Congruent results across laboratories have helped drive further investigation into alternative causes including environmental contaminants and nutritional deficiencies. This work highlights the utility of metagenomic approaches in investigations of emerging diseases and provides a framework for future wildlife mortality events.

**Article Summary Line:** The causative agent of a mass mortality event in passerines remains inconclusive after metagenomic high-throughput sequencing with results prompting further investigation into non-pathogenic causes.

## INTRODUCTION

Wildlife health and diversity are under increasing threats from a multitude of sources, with disease emergence in wildlife having the potential to affect the health of humans and domesticated species (1–3). The development of successful mitigation strategies for emerging infectious diseases in wildlife is often limited by the ability to identify the etiologic agent (4). For example, in May 2015, a mass mortality event was observed in central Kazakhstan in which over half of all saiga antelopes (*Saiga tatarica*) were lost prior to the identification of the etiologic agent and before any mitigation measures could be implemented (5).

Rapid advances in high-throughput sequencing technologies have seen a rise in the number of genomic approaches being applied in disease investigations alongside more traditional techniques, such as histopathology, bacterial culture, virus isolation, and PCR tests (6,7). One such approach is metagenomic next-generation sequencing (mNGS); a culture-independent untargeted technique that can be used to analyze all nucleic acids (i.e., DNA or RNA) within a biological sample. Untargeted approaches, such as mNGS, are unbiased when it comes to capturing all the microbes within a clinical sample, as the majority of microbes can be identified in the absence of a priori assumption (8). This ability is an advantage particularly when the etiologic agent is unknown, and untargeted approaches are increasingly being used to identify pathogenic agents in disease outbreaks affecting humans and livestock (9–11), although it remains relatively uncommon in wildlife (12–15).

Here we highlight the recent use of mNGS to investigate a wildlife mortality event that began in late May 2021 when reports of sick and dead birds were received across the eastern USA. The majority of reports involved nestling and juvenile passerine species including the Common Grackle (*Quiscalus quiscula*), Blue Jay (*Cyanocitta cristata*), European Starling (*Sturnus vulgaris*), American Robin (*Turdus migratorius*), and Northern Cardinal (*Cardinalis cardinalis*), as well as limited reports of non-passeriform avian species that presented with similar clinical signs (i.e., swollen conjunctiva, crusty ocular discharge, head tilt, ataxia, hind limb paresis, and nystagmus). Several diagnostic laboratories launched investigations focused on identifying an etiologic agent using common diagnostic techniques from across multiple disciplines including pathology, virology, microbiology, parasitology, and toxicology (16). Findings from these investigations failed to identify a causative agent but were able to rule out common pathogens and toxicants previously associated with mass avian mortality, including *Salmonella* spp*., Chlamydia* spp., avian influenza viruses, West Nile virus, herpesvirus, *Trichomonas* spp., coccidiosis, and numerous pesticides (17). Several *Mycoplasma* spp. were detected in diseased conjunctiva of some affected birds (unpub. data), but detections were inconsistent, and these bacterial species are commonly detected in non-diseased birds (18).

To further investigate this event, three diagnostic laboratories, namely University of New Hampshire (UNH) Veterinary Diagnostic Laboratory in collaboration with Hubbard Center for Genome Studies, University of Pennsylvania’s Wildlife Futures Program (WFP) in collaboration with Chan Zuckerberg Biohub San Francisco (CZ Biohub SF), and the Infectious Disease Institute at Ohio State University (IDI) undertook mNGS approaches to assist in the detection of a causative agent. Similar mNGS approaches have been previously used in response to mortality events in wild and captive avian species (19–21). Here, we describe the mNGS approaches undertaken by each lab and demonstrate how concurrent approaches can be helpful when investigating the primary cause of a mass mortality event in wildlife.

## METHODS

### Sample collection and processing

During the 2021 mass mortality event in passerines, three labs independently collected samples for mNGS (Figure 1). To summarize, WFP in collaboration with CZ Biohub SF collected whole eye (including conjunctiva) and brain samples from 94 birds including 86 suspected cases and 8 controls in addition to lung, cloacal bursa, and heart blood from 28 birds (20 cases and 8 controls). All suspect cases were selected based on the presence of swollen conjunctiva, eye lesions, and/or crusty ocular discharge. For the WFP samples, suspect cases were all fledglings belonging to one of five species (American Robin, Blue Jay, Common Grackle, Northern Cardinal, and Northern Mockingbird (*Mimus polyglottos*)). Control birds of the same species were sourced from rehabilitation centers in the months following the mortality event (from September to November 2021) where they presented in good nutritional condition with no clinical signs of illness and had died or were euthanized due to acute traumatic injuries (i.e., vehicle collisions or window strikes).

**Figure 1.**
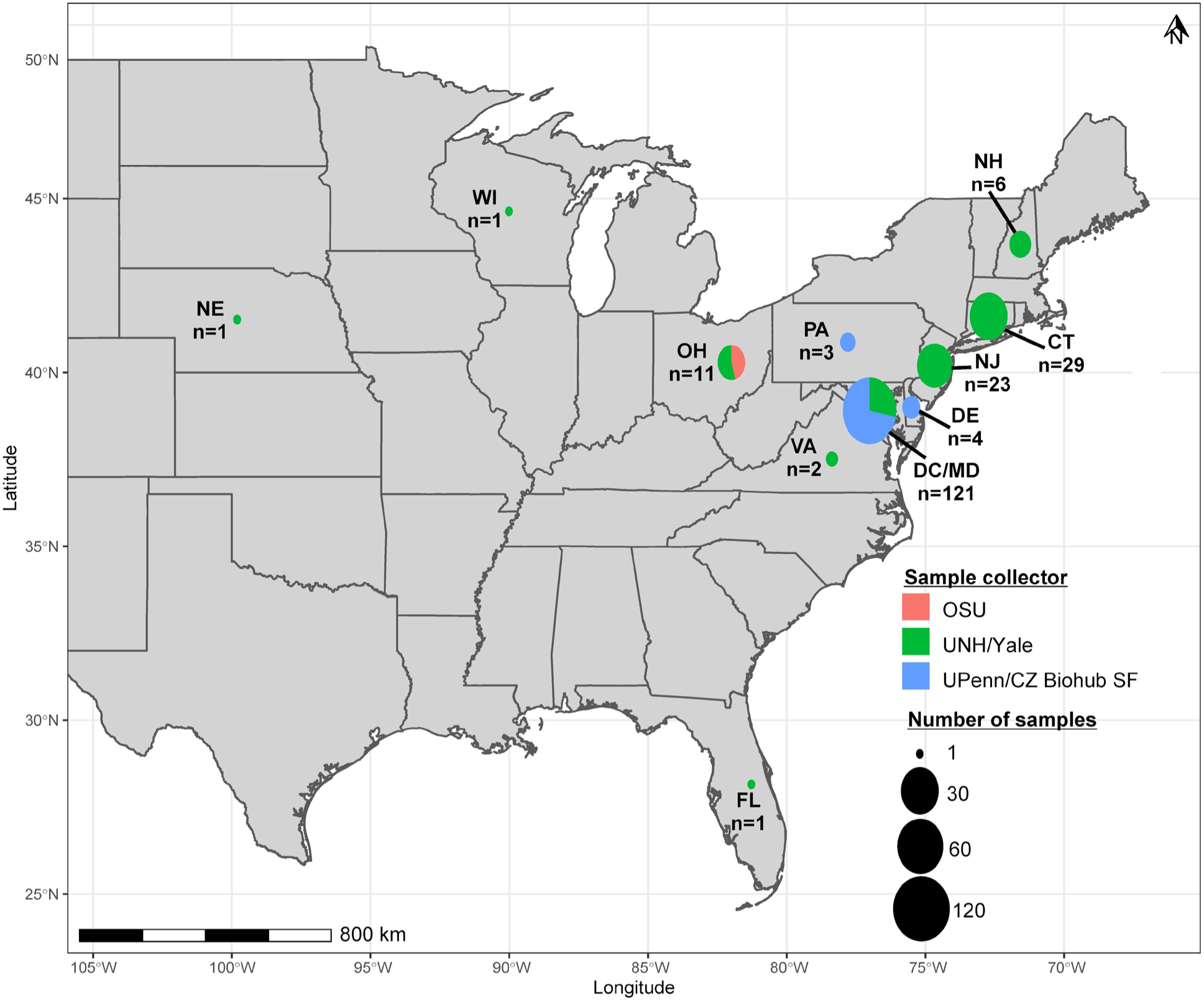
Map showing the distribution of sampled birds (both cases and controls) that were included in the mNGS analyses by state (n=number of birds). The size of the pie chart is proportional to the number of samples while the color of the pie chart indicates the laboratory to which the samples were sent. UPenn: University of Pennsylvania, CZ Biohub SF: Chan Zuckerberg Biohub San Francisco, OSU: Ohio State University, UNH: University of New Hampshire, Yale: Yale University.

In comparison, researchers at UNH collected conjunctiva and ear canal tissue from 103 birds (all suspected cases). These birds were a mixture of adults and fledglings submitted by a number of collaborators including Yale University for which 14 species were identified (American Robin, Blue Jay, Common Grackle, Northern Cardinal, Eastern Phoebe (*Sayornis phoebe*), Rose-breasted Grosbeak (*Pheucticus ludovicianu*s), Rusty Blackbird (*Euphagus carolinus*), European Starling, House Finch (*Haemorhous mexicanus*), Tufted Titmouse (*Baeolophus bicolor*), Cooper’s Hawk (*Accipiter cooperii*), Eastern Screech Owl (*Megascops asio*), Sharp-shinned Hawk (*Accipiter striatus*), and Mourning Dove (*Zenaida macroura*)) in addition to birds that were not identified to species level but were known to belong to one of the following broader taxonomic groups: finch (*Fringillidae*), pigeon/dove (*Columbidae*), thrush (*Turdidae*), sparrow (*Passeridae*), crow (*Corvidae*), or blackbird (*Icteridae*). The majority of these birds were found dead with only a small number having been euthanized at wildlife rehabilitation facilities. Birds outside the passeriform order were treated as suspect cases using the same criteria as suspect cases within the passeriform order (i.e., swollen conjunctiva, eye lesions, and/or crusty ocular discharge).

Lastly, IDI collected brain tissue from 4 birds (all suspected cases). These birds were a mixture of adults and fledglings belonging to one of four species (American Robin, Blue Jay, House Sparrow, and Mourning Dove). All birds were found alive with common clinical signs (swollen conjunctiva, ocular exudate, crusty eyes, and ataxia), and either died during transport or were euthanized at wildlife rehabilitation facilities. Additional details regarding sample collection and processing by each lab are provided in the Appendix.

### Metagenomic next-generation sequencing

At each diagnostic laboratory, different metagenomic approaches were used for extraction, library preparation, and sequencing as summarized in Table 1. Further details are also provided in the Appendix. Some of the major differences between the approaches included the use of different sample types and the extraction of RNA (by WFP/CZ Biohub SF), DNA (by UNH/Yale), or both (by IDI). Following sequencing, the mNGS bioinformatic analysis across the laboratories remained the same with each utilizing the CZ ID metagenomic pipeline - an open-source sequencing analysis platform for identifying microbial sequences within a metagenomic dataset (http://czid.org, *v*6.8). The pipeline removes the avian and human host using STAR (22) and Bowtie2 (23), trims adapters using Trimmomatic (24), filters low-quality reads using PriceSeq (25), filters low-complexity sequences using LZW and identifies duplicate reads using czid-dedup (https://github.com/chanzuckerberg/czid-dedup). The remaining reads are queried on the NCBI nucleotide (NT) and non-redundant protein (NR) databases utilizing GSNAP-L (26) and RAPSearch2 (27), respectively, to determine the microbes (28).

**Table 1.**
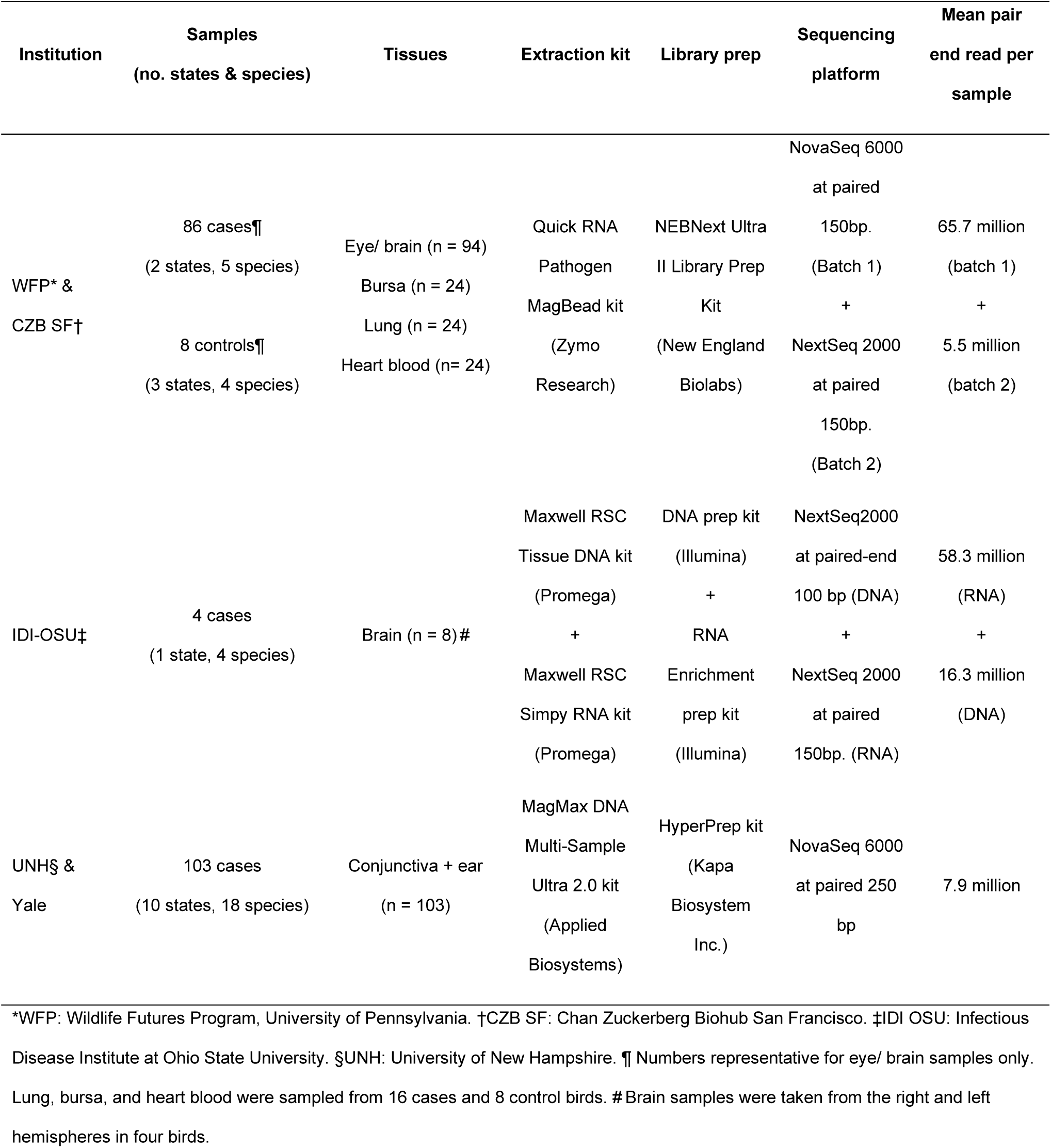
Metagenomic next-generation sequencing (mNGS) approaches taken across three diagnostic laboratories in response to the 2021 mass mortality event in passerines

To account for background contamination, 24 water controls (used when sequencing the WFP/CZ Biohub SF samples) were selected on CZ ID to create a mass-normalized background model for processing their respective samples. Significant microbial hits were called from the normalized unique reads per million (rPM) that mapped to specific species and genera which passed the following threshold filters: z-score ≥ 1 (to denote significant presence in the sample compared to water background), NT rPM ≥ 10 (i.e., ≥10 nucleotide reads per million mapping to specific taxa), NR rPM ≥ 5 (i.e., ≥5 protein reads per million mapping to specific taxa) and average base pair nucleotide alignment ≥ 50 base pairs (i.e., ≥50 average nucleotide reads alignment mapping to specific taxa). To further increase the validity of the microbial hits, select samples were run on the CZ ID Consensus Genome pipeline (*v*3.4.7), to assess the genomic coverage and ensure the number of reads was adequate to obtain consensus genomes. An example of this analysis looking at the validity of West Nile virus detected in a single bird is provided in the Appendix.

### Microbial composition analysis

To investigate the microbial composition, samples were first filtered to remove background taxa (i.e., those present in water controls) by eliminating taxa with a z-score ≤ 1. The proportion of microbes belonging to the taxonomic categories: archaea, bacteria, eukaryotes, or viruses, were reported per respective sample in addition to the two most commonly detected taxa per sample. Further analyses were conducted across the WFP/CZ Biohub SF samples to evaluate differences between the cases and controls. For these analyses, sample reports containing taxonomic relative abundance data for all samples were downloaded from CZ ID and imported into R statistical software (*v*4.2.1; (29)). To investigate the difference in the abundance (NT rPM) of microbes in the eye and brain between the cases and the control group, a Wilcoxon rank-sum test was performed. The significantly differentially abundant microbe genera with *p*-values <0.01 were reported between the groups. Alpha (Simpsons) and beta (Bray Curtis) diversity measures were also calculated using the R package vegan (*v*2.5; (30)) to further investigate the microbial diversity in the eye and brain samples both within and between the case and control groups. The statistical significance in the alpha and beta diversity metrics were evaluated using a Mann-Whitney U test and Permutational Multivariate Analysis of Variance (PERMANOVA) analysis, respectively.

### Data availability

The SRA files of non-host reads for the WFP/CZ Biohub SF and UNH/Yale samples have been deposited with links to BioProject accession numbers PRJNA909835 and PRJNA961153, respectively, in the NCBI BioProject database (https://www.ncbi.nlm.nih.gov/bioproject/).

## RESULTS

### Investigating the microbial composition

No single pathogenic microbe was identified across all the cases, with the most commonly detected microbes varying across diagnostic/research laboratories (Table 2). The species-level distribution consisted mainly of bacterial microbes in both cases and controls with the post-filtering species-level distribution ranging from 43.76% to 95.67% bacterial (mean = 73.08%) across all the samples (Appendix Figures 1,2,3). In addition to the most commonly detected microbes, the presence of other microbes known to be pathogenic to avian species also varied across laboratories. For instance, across the WFP/CZ Biohub SF eye and brain samples, *Avibacterium* spp. (including *A. paragallinarum, A. endocarditidis, A. volantium,* and *A. avium*) and *Mycoplasma* spp. (including *M. gallisepticum, M. pneumoniae,* and *M. mycoides*) were detected in a large proportion of the cases (72.1% and 57.0% respectively) and controls (25.0% and 12.5% respectively), while both *Avibacterium spp.* and *Mycoplasma spp.* were not detected in any of the IDI cases and only in a small proportion from UNH/Yale (21.4% and 6.8% respectively).

**Figure 2.**
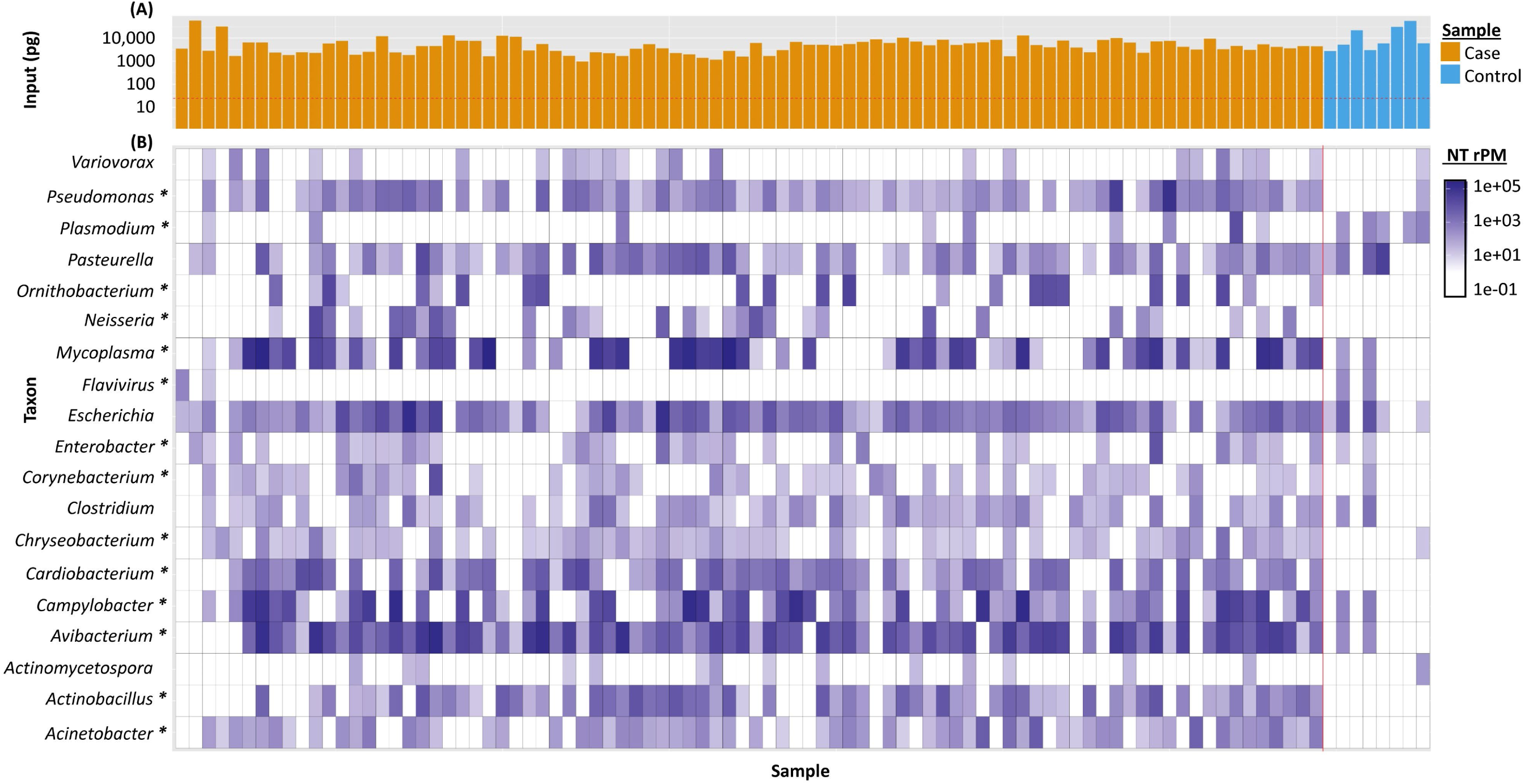
Microbes detected by mNGS in eye and brain samples sorted by genus. Panel (A) denotes sample input (pg) in each sample, whilst the color denotes if the sample was derived from a suspected case (orange) or control (blue). Panel (B) denotes each microbe genus. The asterisk (*) denotes significantly different (p<0.01) microbe presence between the cases (low + high) and controls detected in the sample post-filtering.

**Figure 3.**
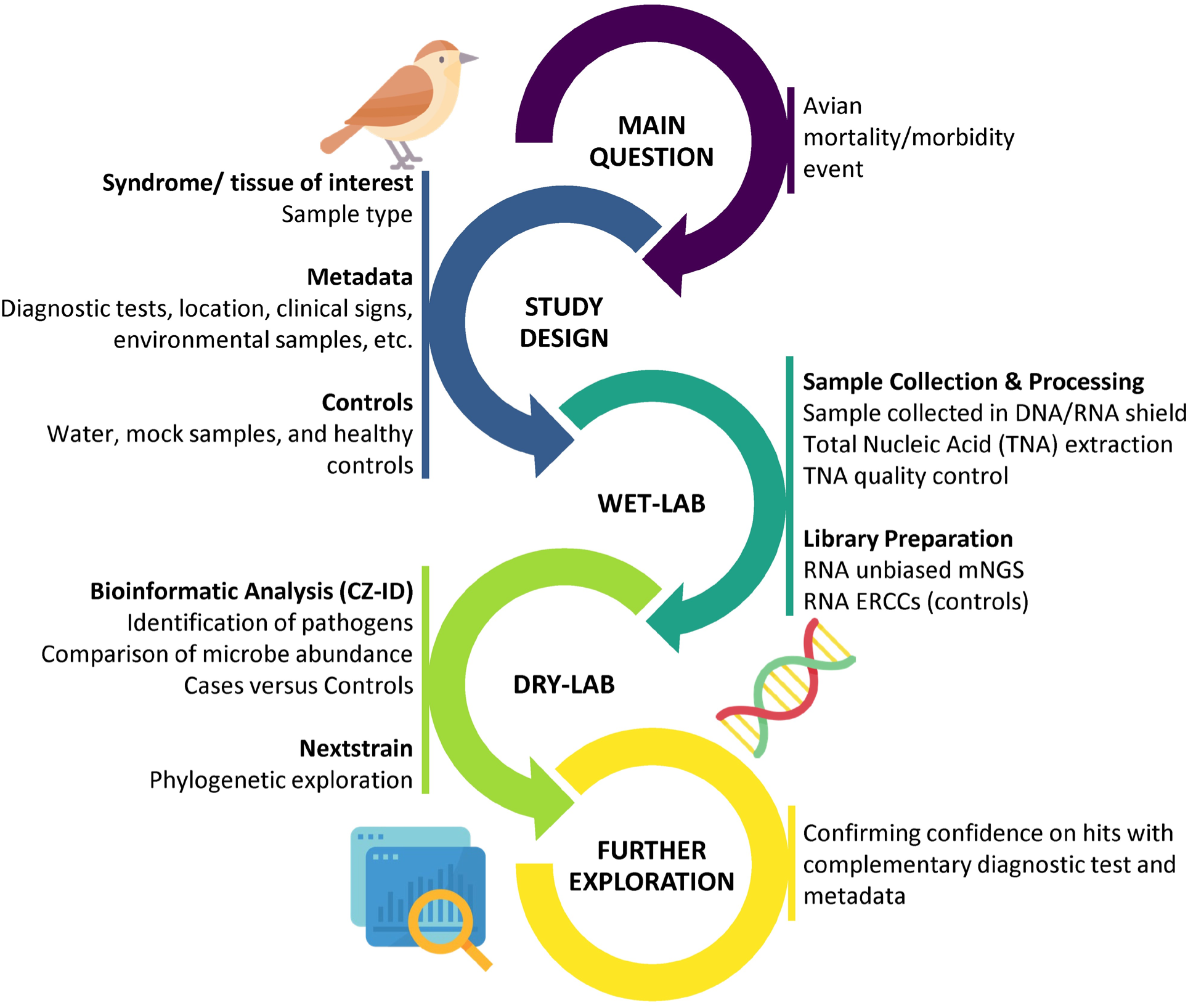
Potential framework for metagenomic next-generation sequencing (mNGS) in wild avian species.

**Table 2.**
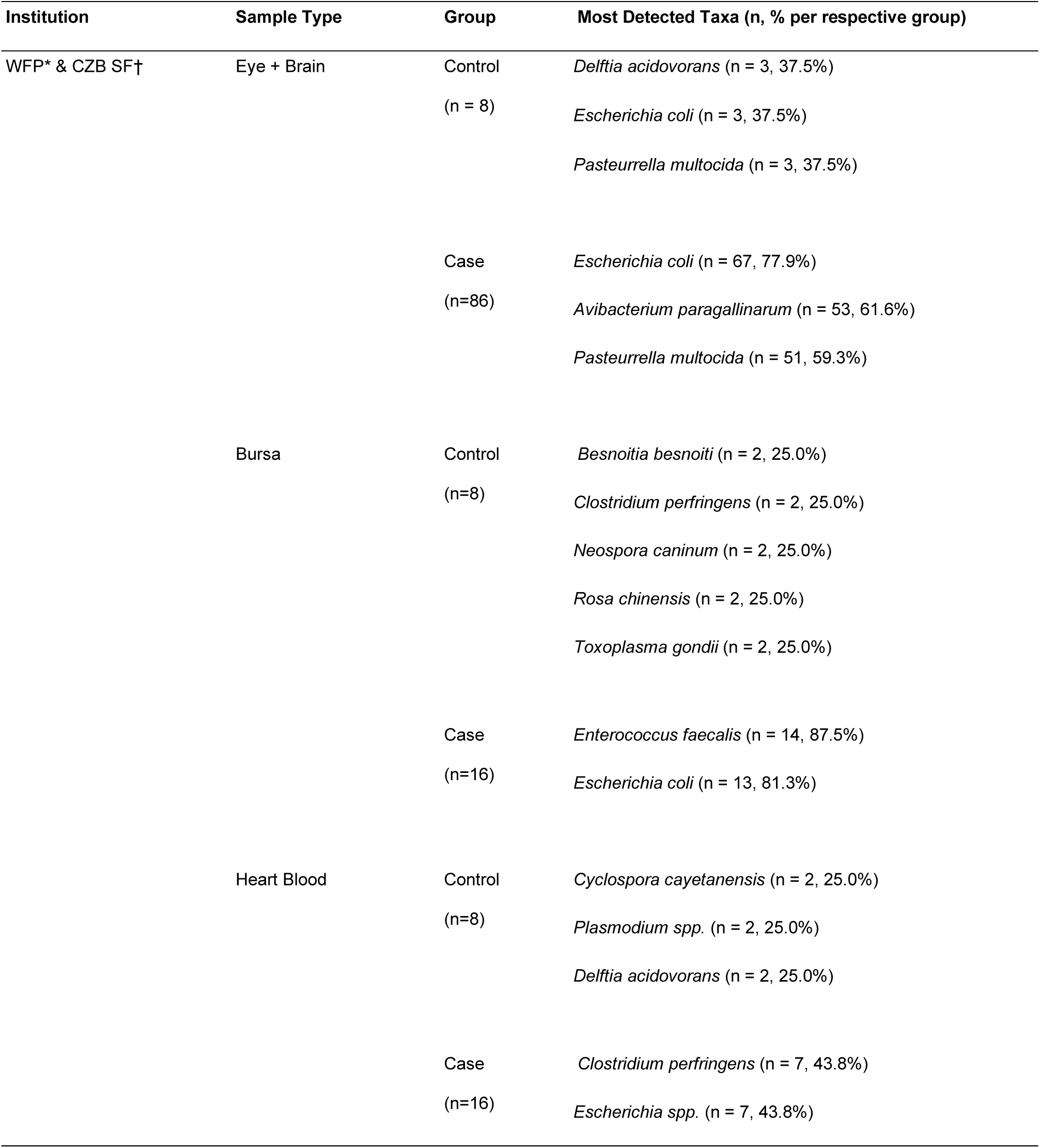
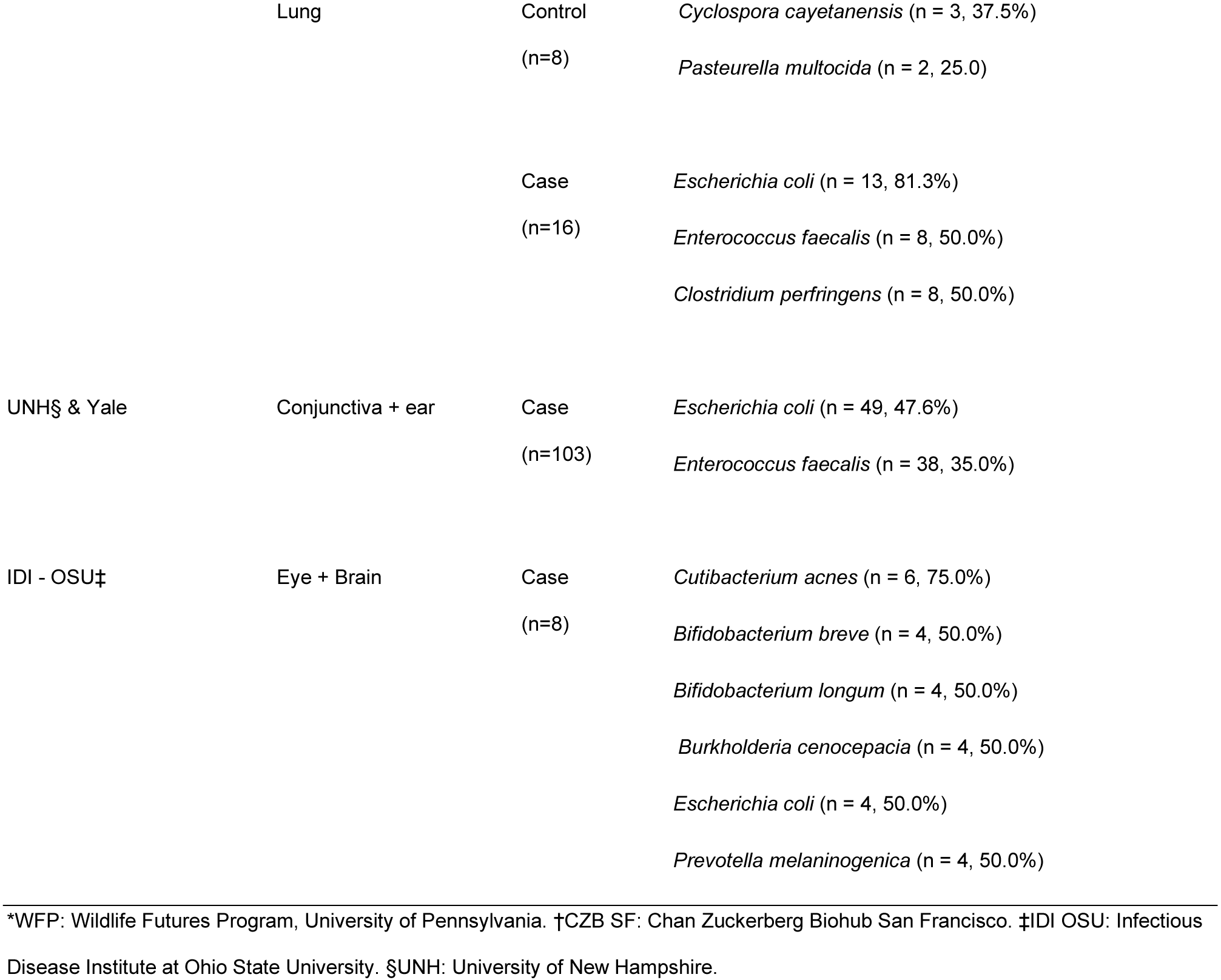
Most frequently detected species taxa per respective sample type and status (i.e., case vs control) following metagenomic next-generation sequencing (mNGS) bioinformatic analysis using the CZ ID metagenomic pipeline (*v*6.8)

*Plasmodium* spp. were also detected in 20.4% of all samples collected (9.7% cases and 65.6% controls) by WFP/CZ Biohub SF but only in a limited number of samples across the other laboratories (Table 3). In comparison, other microbes were detected in the UNH/Yale and IDI samples that were not found in the WFP/CZ Biohub SF; for instance, canarypox virus was detected in 7.62% and 22.2% of the samples, respectively. Furthermore, a comparison of the microbes detected in DNA or RNA libraries at IDI revealed that *Burkholderia cenocepacia*, *Bifidobacterium* spp., and *Prevotella melaninogenica* were only detected in DNA libraries, whilst *Cutibacterium acnes* and *E. coli* were detected in both DNA and RNA libraries.

**Table 3.** *Plasmodium spp*. and *P. relictum* detection across birds following metagenomic next-generation sequencing (mNGS) bioinformatic analysis using the CZ ID metagenomic pipeline (*v*6.8)

| Institution | Sample Type | Group | Microbe |  |
| --- | --- | --- | --- | --- |
|  |  |  | *Plasmodium spp.* | *P. relictum* |
| WFP* & CZB SF† | Eye + Brain | Control | 4 | 0 |
|  |  | Case | 7 | 1 |
|  | Bursa | Control | 2 | 0 |
|  |  | Case | 4 | 1 |
|  | Heart Blood | Control | 6 | 2 |
|  |  | Case | 2 | 2 |
|  | Lung | Control | 7 | 1 |
|  |  | Case | 2 | 1 |
|  | Total Samples |  | 34 | 8 |
|  | Total Birds |  | 15 | 5 |
| UNH‡ & Yale | Conjunctiva + ear | Case | 1 | 0 |
|  |  | Total Samples | 1 | 0 |
|  |  | Total Birds | 1 | 0 |
| IDI - OSU‡ | Eye + Brain | Case | 0 | 0 |
|  |  | Total Samples | 0 | 0 |
|  |  | Total Birds | 0 | 0 |

\*WFP: Wildlife Futures Program, University of Pennsylvania. †CZB SF: Chan Zuckerberg Biohub San Francisco. ‡IDI OSU: Infectious Disease Institute at Ohio State University. §UNH: University of New Hampshire.

Differences in the microbial taxa between case and control samples of the eye and brain were also detected in the WFP/CZ Biohub SF samples. Specifically, the microbial genera of *Mycoplasma* spp*., Campylobacter* spp., and *Avibacterium* spp. were detected at significantly higher levels (*p*-values <0.01) in cases (Figure 2, Table 4). In the diversity analyses, we observed marginally higher, though insignificant differences in Simpson’s alpha diversity (*p*-value = 0.09) species richness in cases versus controls. Meanwhile, the Bray-Curtis beta diversity was significantly different between cases and controls (*p*-values <0.001), suggesting distinct microbial profiles across the two groups.

**Table 4.** Wilcoxon rank test values between cases and controls utilizing NT rPM values from eye/brain samples collected by the Wildlife Futures Program, University of Pennsylvania, and Chan Zuckerberg Biohub

| **Microbe Genera** | **p-value** |
| --- | --- |
| *Acinetobacter* * | 9.38E-06 |
| *Actinobacillus* * | 5.78E-05 |
| *Actinomycetospora* | 1.14E-05 |
| *Avibacterium* * | 8.04E-05 |
| *Campylobacter* * | 4.15E-03 |
| *Cardiobacterium* * | 6.51E-05 |
| *Chryseobacterium* * | 9.78E-05 |
| *Clostridium* | 3.88E-02 |
| *Corynebacterium* | 1.55E-03 |
| *Enterobacter* * | 9.09E-03 |
| *Escherichia* | 2.16E-02 |
| *Flavivirus* * | 2.75E-06 |
| *Mycoplasma* * | 1.14E-03 |
| *Neisseria* * | 8.32E-04 |
| *Ornithobacterium* * | 1.09E-04 |
| *Pasteurella* | 5.83E-01 |
| *Plasmodium* * | 4.16E-04 |
| *Pseudomonas* * | 8.28E-05 |
| *Variovorax* | 1.06E-02 |
| * p-values <0.01 |  |

## DISCUSSION

Here, we describe the metagenomic approaches used to investigate the presence of potential etiologic agent(s) responsible for a mass mortality event in passerines in the eastern USA. After concurrent investigations across multiple diagnostic labs using an array of both targeted (17) and untargeted approaches, no singular pathogenic microbe was identified that would account for the observed morbidity and mortality in these birds, and to date, the results remain inconclusive.

Given the rapid onset and short time period of the event, it is likely that if a pathogen was the primary driver, it would have been detected in a larger percentage of samples. Variations in detection rates due to factors such as the disease stage at which the cases presented, and the tissues collected for sampling are unlikely sufficient to explain the lack of detection of a consistent pathogen across samples. Due to the unpredictable nature of wildlife health events and the reliance on opportunistic sampling, particularly early in the event, some of these factors are easier to control than others. For example, in multi-species events, such as the one reported here, some species may be overrepresented if they thrive in urbanized areas close to humans who can observe the event. The lack of observations from remote regions often results in geographical biases in the samples (31). Temporal biases are also likely, with samples coming from multiple sources and consisting of both morbid animals and animals that were dead for some time, often rendering them suboptimal for diagnostic purposes and raising concerns when comparing samples. Other factors are more easily controlled, such as the tissues selected for sampling, how those tissues are processed, and what diagnostic tests are performed. In this study, whole carcasses and/or tissues of birds suspected to be involved in the outbreak were evaluated at numerous veterinary and wildlife diagnostic laboratories with the utilization of cross-disciplinary diagnostic techniques including pathology, virology, microbiology, parasitology, and toxicology (16). The tissues collected and protocols used by each group varied according to a multitude of factors including resource limitations, time constraints, and funding. This highlights a need for a minimum set of standards to help guide wildlife investigations and ensure some level of consistency across different working groups. A unified framework would also help facilitate collaboration in large multi-state events.

Employing the use of unbiased mNGS for this type of investigation has previously revealed many pathogenic agents including novel pathogens (32–34). Metagenomic findings in this study identified several bacterial pathogens significantly more in cases compared to controls; however, these were deemed unlikely drivers of the mortality event. Foremost, none appeared in a large percentage of samples across groups. In addition, characteristics of the pathology of the specific bacteria identified were inconsistent across birds and with the observed clinical signs. Nevertheless, given the uniqueness of the event, care must be taken in the interpretation of the bacterial pathogens detected. For instance, *Mycoplasma* spp. were found at significantly higher levels in eye and brain samples in cases versus controls. This result could explain the conjunctivitis reported on examination (35); however, *Mycoplasma* spp. are also a common commensal bacterium in many avian species (18). Thus, it is possible the detection of this microbe was due to opportunistic and/or subclinical infections rather than being a primary causative agent - particularly given it was still found in a large percentage of control birds. In addition, *Mycoplasma* spp. were not consistently found across laboratories. Together, this suggests *Mycoplasma* spp. was not primarily responsible for the mortality event.

*Avibacterium* spp. were also detected at significantly higher levels in the cases than the controls; however, the clinical signs characteristic of *Avibacterium* spp. infections are more consistent with respiratory disease (i.e., mouth breathing, swollen sinuses, and nasal discharge (36,37), which were not observed. However, these clinical signs are typically described for chickens (*Gallus gallus domesticus*), with infections in wild avian species rarely reported and not well described; therefore, we cannot be certain that wild avian species would experience the same clinical signs. For example, a recent study reported severe periocular swelling, periocular skin crusting, fibrinous sinusitis, and conjunctivitis in wild turkeys infected with a novel clade of *Avibacterium* (38). The presence of multiple pathogenic agents also makes it difficult to determine the contribution of different microbes. For instance, 45.3% of cases for which both eye and brain were assessed were coinfected with *Mycoplasma* and *Avibacterium* spp. This coinfection has also not been well described and thus, the resulting clinical signs are unknown; however, without more control cases or experimental infection trials, it is difficult to draw further conclusions. Yet, despite the limited number of controls, the beta diversity analysis did reveal a significant difference in the microbial compositions between the clinical cases and controls, suggesting that the former contained a different microbial profile which could have contributed to the morbidity and mortality.

Although no causative agent for the mortality event was identified, findings highlight the potential of using non-targeted approaches, such as mNGS, to help describe the microbial community circulating in wild populations including both pathogenic and non-pathogenic microbes (39,40). For example, the detection of *Avibacterium paragallinarum* in this study is important to document as it is known primarily as a respiratory disease affecting chickens and is rarely described to cause disease in wild avian species (41). Generating baseline data and understanding how it changes over time further enables downstream comparisons to be made between diseased and healthy individuals and supports diagnostic responses to future mortality events. For example, the finding of *Mycoplasma* spp. in the control birds highlights the importance of using controls to help elucidate potential causes of disease in wildlife (42,43). Further, with the added health challenge posed by continual changes in the environments in which these birds live (e.g., landscape, climate, accumulation of potentially toxic substances), baseline microbes could be used as an indicator of the changing health status of an animal population (44).

Though mNGS is a powerful tool for detecting potential microbial pathogens, variability in experimental protocols, such as the sampling procedures, processing, and data analyses, can lead to artifacts that may result in incorrect conclusions and false negative detections (45,46). Each of the three laboratories in this study used a different experimental design, which enabled us to compare results and make recommendations on the best practices for conducting an mNGS investigation in response to a wildlife mass mortality event. Here we suggest a framework (Figure 3) to assist in addressing some of the limitations of this study for future explorations.

Firstly, developing a strong case definition is pivotal to help guide sample selection. In the early stages of the investigation, a case definition may be primarily based on the range of clinical signs present in the affected population; however, as more cases are identified it is important to revisit the case definition and include information regarding both clinical, laboratory, and pathologic characteristics as well as information on the affected individuals (e.g., species, age, etc.), and any geographical or temporal characteristics. A strong case definition also helps in the selection of high-quality controls from the same population as the cases to provide background microbiota data from healthy specimens when performing downstream comparative analyses. Controls should only be selected if their cause of death is known and unrelated (e.g., they show no clinical and/or pathological signs of disease and/or are from a different area/population). Water controls during sample extraction should also be implemented to remove background contamination during the library preparation, as well as mock samples that may be used to spot contamination that may occur during sample collection and processing.

During sample collection, utilizing a nucleic acid stabilizer is crucial to assisting with retaining the total nucleic acid integrity and inactivating any infectious agents. We suggest starting with RNA libraries for the initial pass to reveal all actively replicating microbes within the samples that may be contributing to an active infection, while also allowing the capture of RNA viruses. However, if the RNA libraries yield no results, DNA libraries can be performed. For microbe detection in mNGS data, several steps can be taken to help increase the confidence in microbe detection further downstream in the analysis, including the implementation of threshold filters to help validate and increase confidence in the presence of detected pathogens, incorporating controls to compare microbial profiles and abundances between the case and control groups, and utilizing metadata to aid in the logical assessment of potentially pathogenic microbes that have been detected. It is also important to consider that any reference-based assembly could miss novel pathogens currently unavailable in the reference database, and further interrogation of the reads may be required (47,48). Our suggested framework for mNGS approaches (Figure 3) helps ensure that steps have been taken to minimize artifacts in the data. Here, parallel analyses across three diagnostic laboratories revealed no single pathogen associated with the 2021 mass mortality event in passerines. Findings suggest that the underlying mortality is not due to pathogenic microorganisms and have guided the investigation to refocus time and resources on other potential factors, such as dietary deficiencies, to explain the mortality event.

## ACKNOWLEDGMENTS

We are grateful to all wildlife rehabilitators, biologists, veterinarians and veterinary staff, volunteers, members of the public, and responding state wildlife agencies who assisted with sample collection. We also want to extend our thanks to the CZ Biohub SF Genomics team for the sequencing support, the Wildlife Futures team for sampling assistance, and the faculty and staff at the Pennsylvania Animal Diagnostic Laboratory System-New Bolton Center who contributed to the sample collection, diagnostics, and shipping. S.S.G. was supported by the Robert J. Kleberg, Jr. and Helen C. Kleberg Foundation, while A. J. A was supported by an NSF Postdoctoral Fellowship in Biology (Award #2010904). UNH sequencing was funded through the USDA NIFA NH00694 awarded to W. K. Thomas.

## ADDRESS FOR CORRESPONDENCE

**Roderick Gagne**, Department of Pathobiology, Wildlife Futures Program, University of Pennsylvania School of Veterinary Medicine, New Bolton Center, 382 W. Street Road, Kennett Square, PA 19348, USA.

