## Appendix for "Metagenomics for pathogen detection during a wildlife mortality event in songbirds"

Chan Zuckerberg Biohub, San Francisco, CA, USA (L. Mwakibete, V. Ahyong, N. Neff, A. M. Detweiler, C.M. Tato)

Chan Zuckerberg Initiative, Redwood City, CA, USA (K. Kalantar)

Department of Pathobiology, Wildlife Futures Program, University of Pennsylvania School of Veterinary Medicine, New Bolton Center, Kennett Square, PA, USA (S. S. Greening, E. Anis, E. A. Miller, L. A. Murphy, J. C. Ellis, R. B. Gagne)

Department of Pathobiology, PADLS New Bolton Center, University of Pennsylvania School of Veterinary Medicine, New Bolton Center, Kennett Square, PA USA (E. Anis, L. A. Murphy)

Department of Ecology and Evolutionary Biology, Yale University, New Haven, CT, USA (C. B. Ogbunugafor, A. J. Ayala)

University of New Hampshire, New Hampshire Veterinary Diagnostic Lab, Durham, NH, USA (D. B. Needle, T. Baillargeon, S. Tanguay)

Hubbard Center for Genome Studies, University of New Hampshire, Durham, NH, USA (J. L. Sevigny, L. M. Gordon, S. D. Simpson, W. K. Thomas)

Infectious Diseases Institute, The Ohio State University, Columbus, OH, USA (M. Oglesbee, S. A. Faith)

Southeastern Cooperative Wildlife Disease Study and Department of Pathology, College of Veterinary Medicine, University of Georgia, Athens, GA, USA (N. M. Nemeth)

Department of Pathology, College of Veterinary Medicine, University of Georgia, Athens, GA, USA (N. M. Nemeth)

### METHODS EXTENSION

#### *Wildlife Futures Program, Pennsylvania Animal Diagnostic Laboratory System-New Bolton Center and Chan Zuckerberg Biohub San Francisco*

Samples were processed in two batches. 82 birds were collected by Second Chance Wildlife Center in Gaithersburg, MD between 22 May and 26 June 2021. These birds were all fledglings belonging to one of five species (American robin, blue jay, common grackle, northern cardinal, and northern mockingbird) that presented with eye lesions. All the birds were either euthanized on arrival using an overdose of isoflurane inhalant anesthesia or arrived deceased. Whole birds were double bagged in resealable plastic bags and stored at -18 °C. For the first batch, eye and brain samples were collected from the 82 frozen birds with all necropsies carried out in biosafety cabinets. To collect eye tissue, dissection scissors were used to excise one ocular globe which was then placed in a cryovial of DNA/RNA Shield (Zymo Research, CA, USA). For brain tissue, the skull was opened at the orbit of the enucleated eye, and a $5mm^3$ section of the cerebrum was removed and placed in the same cryovial. Instruments were disinfected between samples by wiping off grossly visible tissue, then placing them in 15% bleach for 10 minutes followed by ethanol for a further 5 minutes, and then rinsed in distilled water.

For the second batch, additional lung, bursa, and heart blood samples were removed from a subset (n=16) of the 82 birds used in batch one in addition to eight control birds that were submitted to Tri-State Bird Rescue and Research Inc., Newark, DE with a known cause of death (i.e., euthanized due to traumatic injuries) and no clinical signs compatible with the morbidity/mortality event as well as four additional birds believed to be cases. To collect the additional tissue samples, sterile dissection scissors were used to make a ventral midline incision on the coelom giving access to the bursa and the lungs. An incision was then made to the right ventricle to allow frozen blood to be removed from the heart. All samples were placed in separate cryovials with DNA/RNA Shield and transported overnight to CZ Biohub (San Francisco, CA, USA) for sequencing.

During sequence preparation, samples were aliquoted into bead-bashing tubes with at least 400µL of DNA/RNA shield. Samples were bead bashed at 30Hz for 1 min, put on ice for 1 min, and bead bashed again for 1 min at 30Hz. Total nucleic acid (TNA) was extracted from 200µL of bead-bashed samples in 1X shield using the quick-DNA/RNA Pathogen MagBead kit

(Zymo Research). Extracted TNA was DNAsed to attain RNA and ran on a TapeStation for quality control to examine RNA integrity. Negative water controls were used for background contamination as well as a positive control ERCC spike-in (RNA standard dilution series from External RNA Controls Consortium). FastSelect -rRNA HMR (Qiagen) was used for human RNA ribosomal depletion at 1:10X. RNA was reverse transcribed to attain cDNA which was used to construct and barcode sequencing libraries using the NEBNext Ultra II Library Prep Kit (New England Biolabs). The RNA sequencing libraries underwent 150-nt paired-end Illumina sequencing separately per respective batch with Illumina NovaSeq 6000 for batch 1(Illumina) and NextSeq 2000 (Illumina) for batch 2.

***New Hampshire Veterinary Diagnostic Laboratory***

Samples had been collected by a mixture of wildlife agency biologists and wildlife rehabilitation organizations from across 10 states (CT, NJ, DC, NH, OH, MD, VA, WI, NE, FL) between 22 May and 14 July 2021. These birds were a mixture of adults and fledglings for which 14 species were identified (American robin, blue jay, common grackle, northern cardinal, eastern phoebe (*Sayornis phoebe*), rose-breasted grosbeak (*Pheucticus ludovicianus*), rusty blackbird (*Euphagus carolinus*), European starling (*Sturnus vulgaris*), house finch (*Haemorhous mexicanus*), tufted titmouse (*Baeolophus bicolor*), Cooper's hawk (*Accipiter cooperii*), eastern screech owl (*Megascops asio*), sharp-shinned hawk (*Accipiter striatus*), and mourning dove (*Zenaida macroura*)) in addition to birds that were not identified at the species level but were known to belong to one of the following taxonomic categories: finch sp., pigeon/ dove sp., thrush sp., sparrow sp., *Corvus* sp., or blackbird sp. The majority of these birds were found dead with only a small number having been euthanized at wildlife rehabilitation organizations. All birds were stored whole at -18 °C then thawed in batches for tissue collection. For each bird, conjunctiva and ear canal samples were collected. Instruments were disinfected between samples by placing them in Virex for 10 minutes before rinsing them in distilled water. Chicken (*Gallus gallus domesticus)* lungs and intestines were used as controls.

In preparation for sequencing, 20mg of the collected tissue from each bird was added to a fluid mixture containing 20 µL enhancer solution, 500 µL extraction buffer, and 40 µL proteinase K in a 1.5 mL tube. The samples were then incubated for 2-4 hours at 65°C. After the incubation process, 400 µL of the supernatant was transferred to a deep-well plate and the samples were stored at -20°C overnight. The following morning, plates were thawed and then 400 µL of the binding solution, 40 µL DNA binding beads, and 10 µL of RNase A were added to each well. For the extraction process, a KingFisher Flex 96 (Thermo Fisher Scientific™) was used such that plate 1 consisted of 1000 µL of wash solution per well from the MagMax DNA Multi-Sample Ultra 2.0 kit, plate 2 consisted of 1000 µL 80% ethanol per well, plate three had 500 µL 80% ethanol per well, and plate four had 125 µL elution solution (MagMax DNA Multi-Sample Ultra 2.0 kit) per well. After extraction, library preparation was performed using the Kapa HyperPrep kit with TruSeq adapters, and sequencing was completed on an Illumina NovaSeq 6000 instrument to produce 250 bp paired-end reads.

***Infectious Disease Institute at Ohio State University***

Samples were collected by Columbus Zoo and Ohio Wildlife Center between 10 July and 16 July 2021. These birds were a mixture of adults and fledglings belonging to one of four species (American robin, blue jay, house sparrow, and mourning dove). All the birds were found alive with common clinical signs (swollen conjunctiva, ocular exudate, crusty eyes, and ataxia), and either died in transport or were euthanized at the Ohio Wildlife Center using an intracardiac Euthasol® injection. For storage, all birds were decapitated, and their heads were stored in a double ziplock bag at -18°C ready for tissue collection. Before dissection, the 4 selected sample bird heads were thawed at room temperature for 15 minutes. After thawing, complete gross post-mortem examinations were performed before the brain was resected with a brain stem and divided into two hemispheres along the longitudinal fissures with one half used for RNA library prep and the other for DNA library prep. Hemispheres were dissected into smaller pieces, weighed, and placed into individual 50 mL tubes. DNA and RNA were extracted using the MaxWell RSC Tissue DNA (Promega) and Maxwell RSC Simpy RNA (Promega) kits respectively. DNA libraries were built with the DNA Prep kit from Illumina, using 22ng genomic DNA input for each respective library. DNA libraries were sequenced on the NextSeq2000 at paired-end 100 bp. RNA libraries were built with the Illumina RNA Enrichment Prep kit. 50ng RNA input was used for each respective library. Prior to RNA library preparation, FastSelect -rRNA HMR (Qiagen) was used to deplete host-ribosomal RNA at the RT step. RNA libraries were sequenced on the NextSeq 2000 at paired-end 150bp.

#### ***Consensus Genome Pipeline and Phylogenetic Analysis - West Nile Virus***

To further increase the validity of the microbial hits, select samples were run on the CZ ID Consensus Genome pipeline (*v*3.4.7). The consensus genome pipeline aligned non-host reads to the top reference genome identified by CZ ID. Aligned reads were then trimmed (Trim Galore; https://github.com/FelixKrueger/TrimGalore) to remove adapters, low-quality reads (Phred score <20), and short sequences less than 20bp. Consensus bases were called if they had a coverage depth of 5 or more reads. Bases that weren't called were identified as N, and SNPs were called using SAMtools and BCFtools (1). The Nextstrain platform (2) was then utilized to perform phylogenetic inference on microbes of interest using the consensus genomes obtained from this analysis. For instance, West Nile virus was selected for further phylogenetic analysis to investigate how the detected sequences compared to publicly available sequence data from the Virus Pathogen Resource Database (ViPR; (3)) and the WNV Nextstrain build developed by the Grubaugh Lab (4) between April 2015 and December 2021. Nextstrain Augur was then used to perform a multiple sequence alignment of input sequences using *MAFFT* (5) and trimmed it to the reference genome, removing any sequence insertions which introduced gaps in the reference genome. IQ-TREE (6) was used to generate a maximum likelihood genetic divergence tree, followed by TreeTime (7) to temporally resolve the tree which estimates the molecular clock rate within a maximum likelihood framework. The generated tree was exported for visualization in Nextstrain Auspice.

Overall, West Nile virus was detected by mNGS in a single bird across all four sample types (i.e., eye/ brain, bursa, heart blood, and lung) with four full genomes (>99%) recovered from each sample type. The phylogenetic tree containing the 4 consensus genomes alongside other recent West Nile virus genomes can be found in Appendix Figure 4A. This tree includes 216 West Nile virus genomes (including the 4 we generated during this study). Our samples are part of a clade defined by a C6675T mutation. The inferred date for the most recent common ancestor is December 2010 (95% confidence interval (CI): November 2009 - February 2012). The 4 sequences cluster together with 48 additional mutations from the basal virus. The consensus genomes also cluster closely with other WNVs from the USA, specifically with genomes recovered from bird and mosquito hosts (Appendix Figure 4B).

### FIGURES

**Appendix Figure 1:** Overall microbial composition by broad taxonomic categories in **(A)** pre-filtered and **(B)** post-filtered metagenomic sequences obtained from the conjunctiva and ear canal tissue of case birds submitted to the New Hampshire Veterinary Diagnostic Laboratory. Uncategorized reads include vectors, uncultured micro/organisms, uncultured prokaryotes, unidentified soil organisms, and taxa with neither family nor genus classification. (Filters applied: NT rPM ≥ 10, NR rPM ≥ 5, and average NT alignment ≥ 50 base pairs).

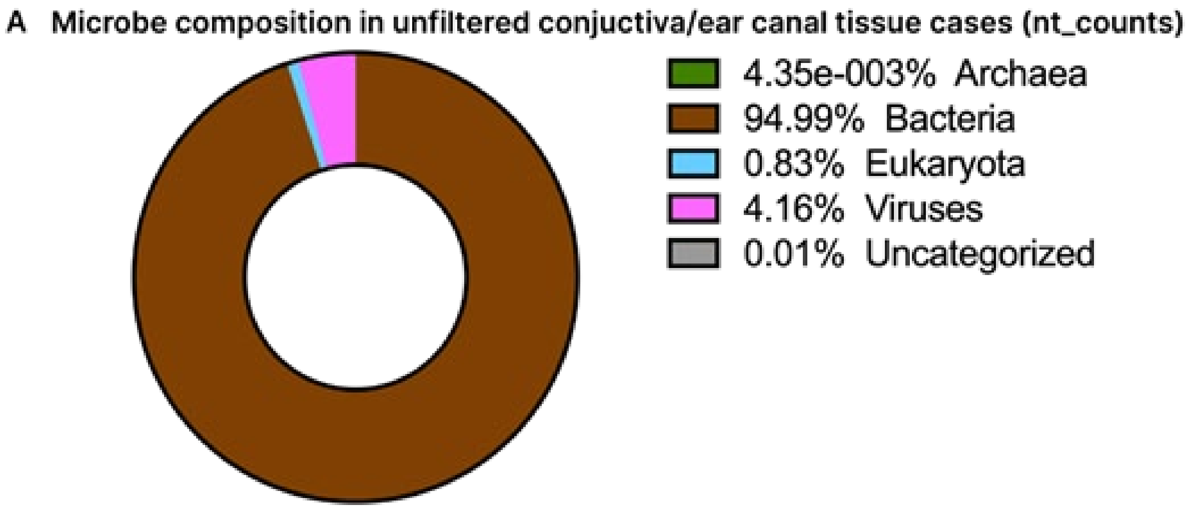


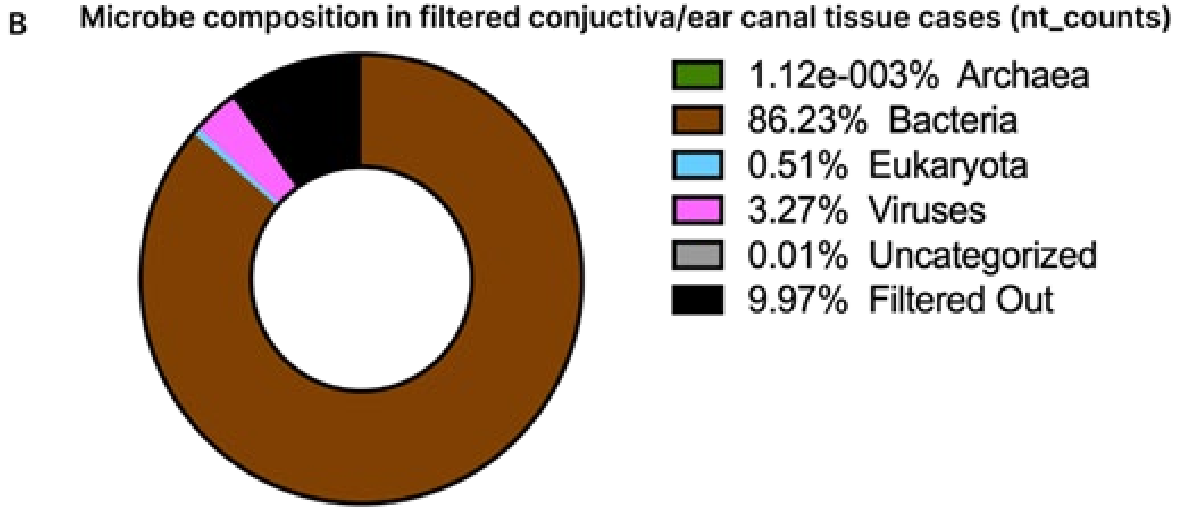

**Appendix Figure 2:** Overall microbial composition by broad taxonomic categories in **(A)** pre-filtered and **(B)** post-filtered metagenomic sequences obtained from the brain tissue of case birds submitted to the Infectious Disease Institute at Ohio State University. Uncategorized reads include vectors, uncultured micro/organisms, uncultured prokaryotes, unidentified soil organisms, and taxa with neither family nor genus classification. (Filters applied: NT rPM ≥ 10, NR rPM ≥ 5, and average NT alignment ≥ 50 base pairs).

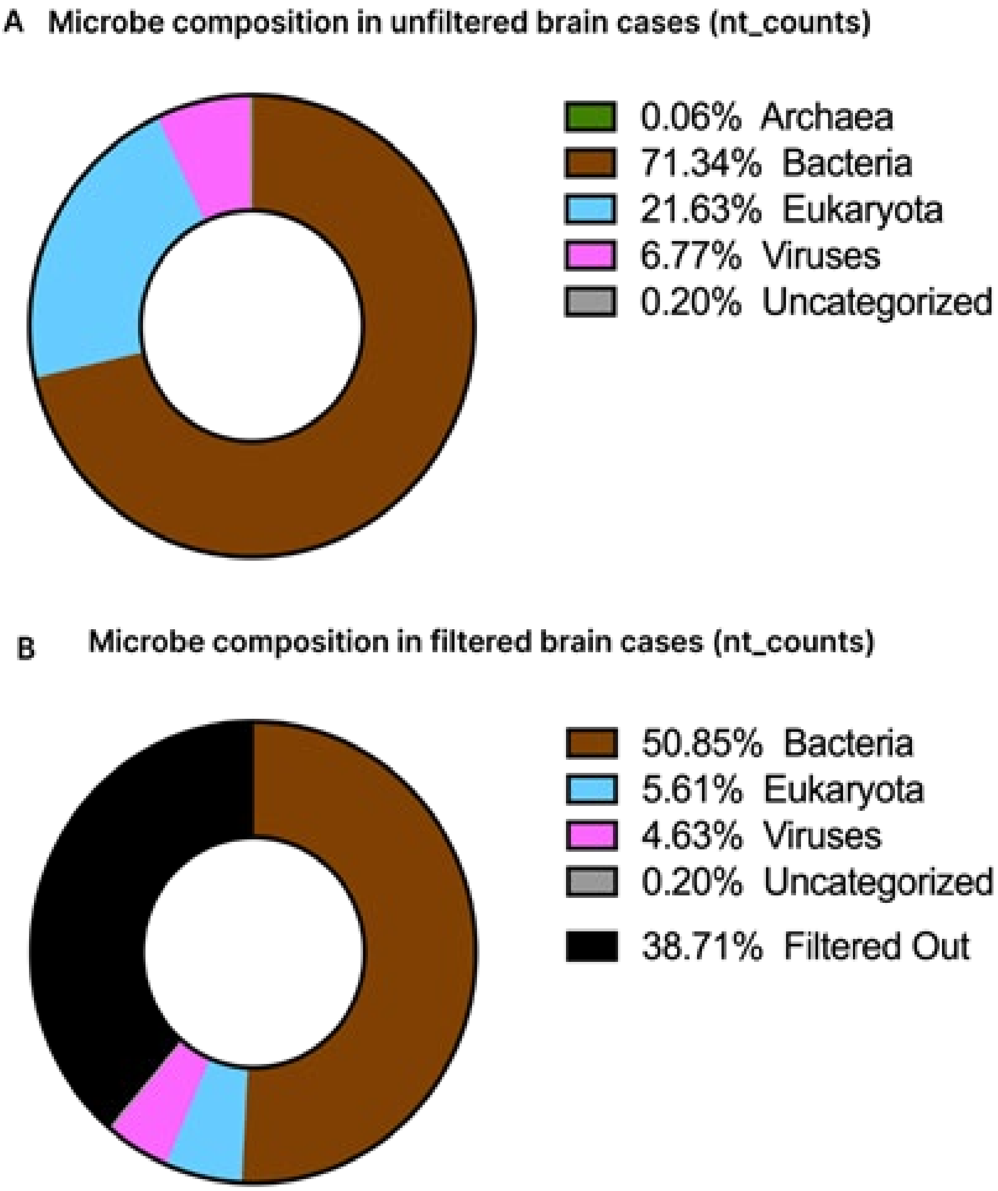

**Appendix Figure 3:** Overall microbial composition by broad taxonomic categories in **(A)** pre-filtered and **(B)** post-filtered metagenomic sequences obtained from the eye and brain tissues of case birds submitted to the Wildlife Futures Program and Chan Zuckerberg Biohub San Francisco and post-filtered metagenomic sequences from **(C)** eye and brain samples from controls, **(D)** cloacal bursa samples from cases, **(E)** cloacal bursa samples from control, **(F)** heart blood from cases, **(G)** heart blood from controls, **(H)** lung samples from cases, and **(I)** lung samples from controls. Uncategorized reads include vectors, uncultured micro/organisms, uncultured prokaryotes, unidentified soil organisms, and taxa with neither family nor genus classification. (Filters applied: NT rPM ≥ 10, NR rPM ≥ 5, and average NT alignment ≥ 50 base pairs).

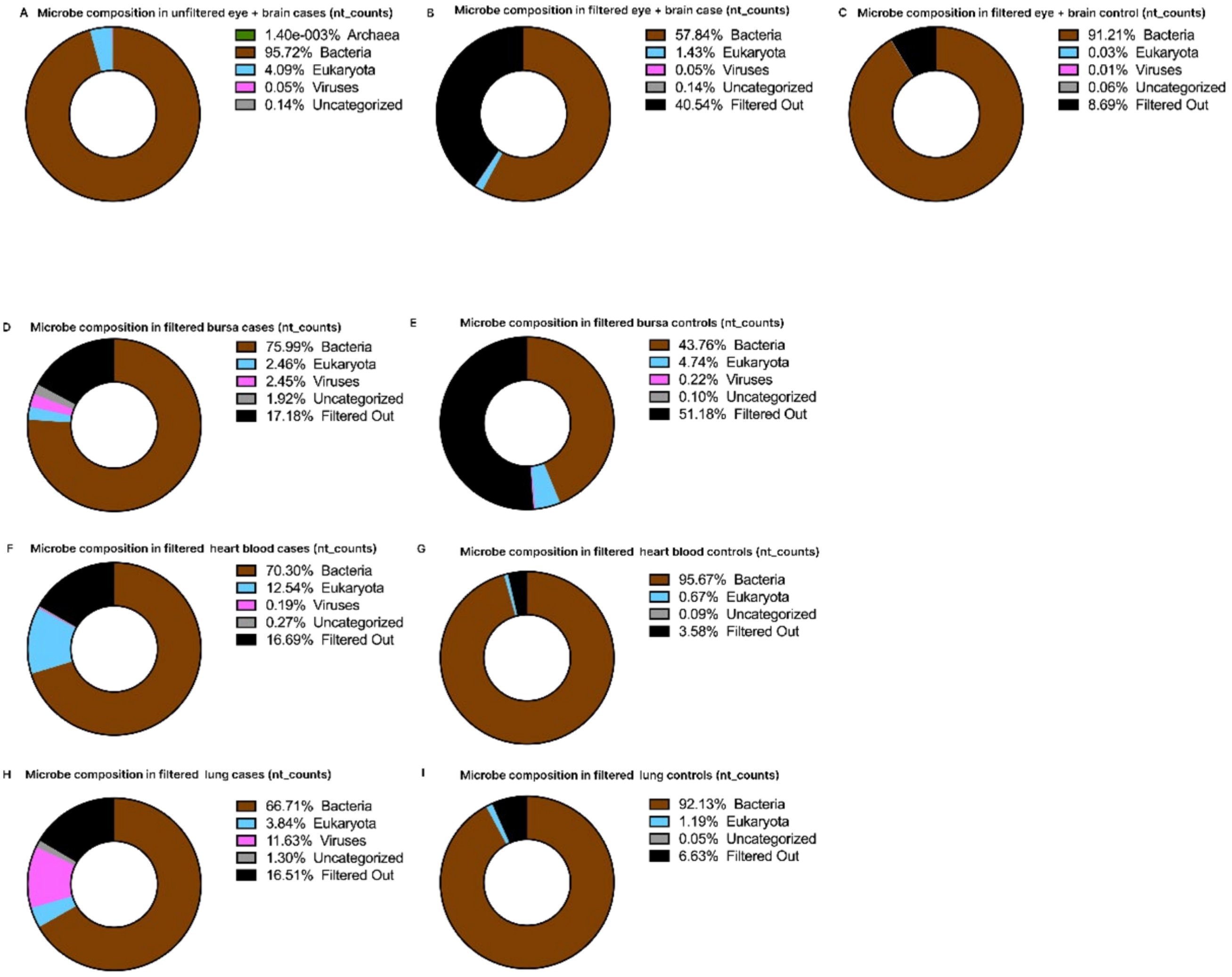

**Appendix Figure 4:** Phylogeographic analysis of West Nile virus (WNV) genomes generated using mNGS. **(A)** Temporally resolved phylogeny of WNV genomes based on 212 West Nile virus genomes from the Virus Pathogen Resource Database (VIPR), the WNV Nextstrain build developed by the Grubaugh Lab (https://github.com/grubaughlab/WNV-nextstrain), and the 4 WNV genomes generated in this study denoted by the black rectangle. **(B)** The same phylogeny as in panel A, with the 4 WNV virus genomes from this study denoted by the black rectangle. In panel A, tip colors indicate the country of origin, whilst, in panel B, tip colors indicate the host. Sub-sampling was used to get 100 sequences per country per year and month.

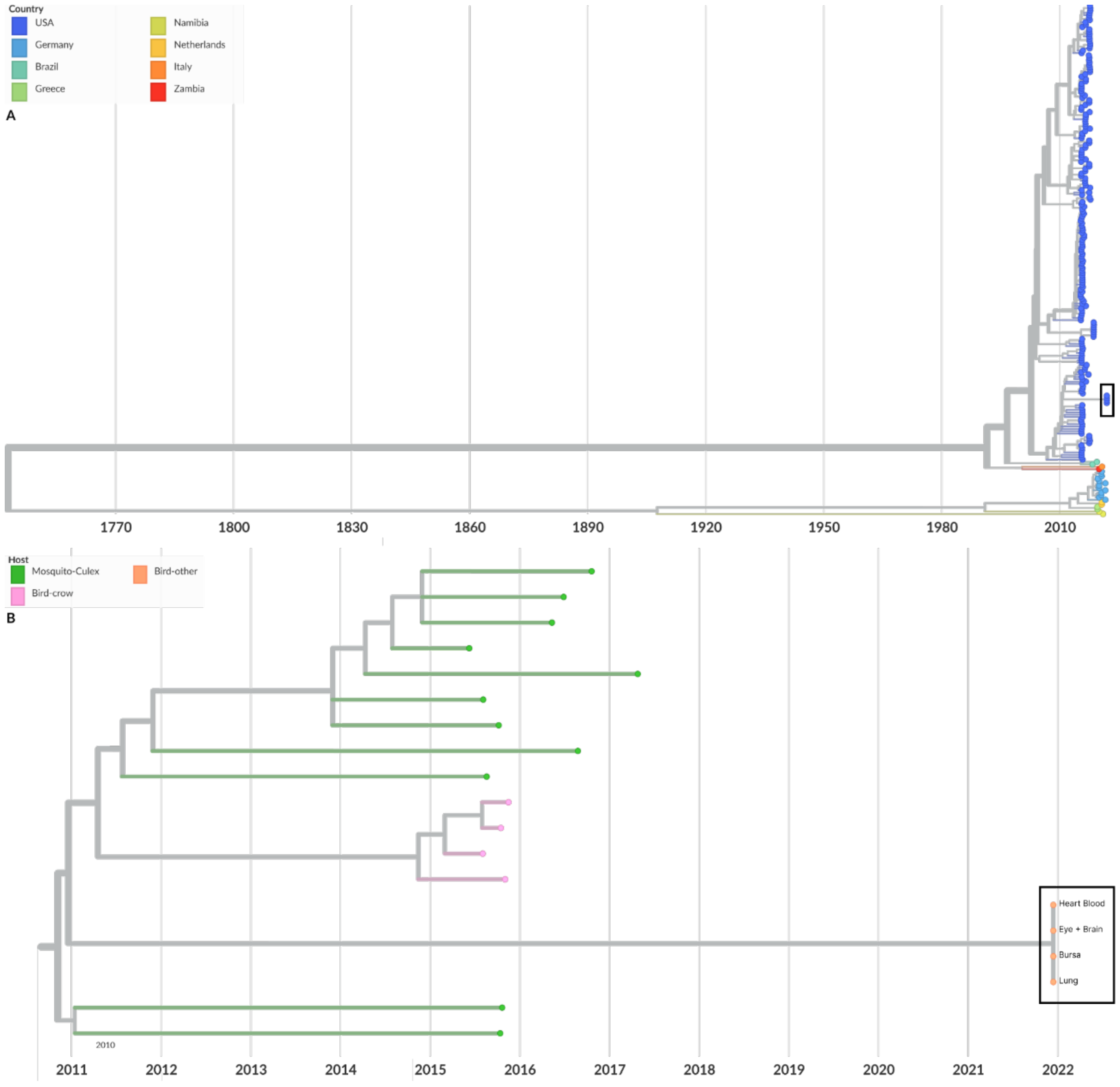
